# Occipital tACS bursts during a visual task impact ongoing neural oscillation power, coherence and LZW complexity

**DOI:** 10.1101/198788

**Authors:** Marta Castellano, David Ibanez-Soria, Eleni Kroupi, Javier Acedo, Michela Campolo, Xenia Martinez, Aureli Soria-Frisch, Josep Valls-Sole, Ajay Verma, Giulio Ruffini

## Abstract

Little is known about the precise neural mechanisms by which tACS affects the human cortex. Current hypothesis suggest that transcranial current stimulation (tCS) can directly enhance ongoing brain oscillations and induce long - lasting effects through the activation of synaptic plasticity mechanisms [1]. Entrainment has been demonstrated in in - vitro studies, but its presence in non-invasive human studies is still under debate [2,3]. Here, we aim to investigate the immediate and short-term effects of tACS bursts on the occipital cortex of participants engaged in a change – of - speed detection task, a task that has previously reported to have a clear physiology - behavior relationship, where trials with faster responses also have increased power in γ - oscillations (50 - 80 Hz) [4]. The dominant brain oscillations related to the visual task are modulated using multichannel tACS at 10 and 70 Hz within occipital cortex. We found that tACS stimulation at 10 Hz (tACS _10_) enhanced both α (8 - 13 Hz) and γ oscillations, in hand with an increase in reaction time (RT) in the change – of - speed detection visual task. On the other hand, tACS at 70Hz desynchronized visual cortices, impairing both phase - locked and endogenous γ - power while increasing RT. While both tACS protocols seem to revert the relationship reported in [4], we argue that tACS produces a shift in attentional resources within visual cortex while leaving unaltered the resources required to conduct the task. This theory is supported by the fact that the correlation between fast RT and high γ- power trials is maintained for tACS sessions too. Finally, we measured cortical excitability by analyzing Event – Related - Potentials (ERP) Lempel – Ziv - Welch Complexity (LZW). In control sessions we observe that lower γ - LZW complexity correlates to faster reaction times. Both metrics are altered by tACS stimulation, as tACS _10_ decreased amplitude of the P300 peak, while increasing γ- LZW complexity. To this end, our study highlights the nonlinear cross - frequency interaction between exogenous stimulation and endogenous brain dynamics, and proposes the use of complexity metrics, as LZW, to characterize excitability patterns of cortical areas in a behaviorally relevant timescale. These insights will hopefully contribute to the design of adaptive and personalized tACS protocols where cortical excitability can be characterized through complexity metrics.

**Additional Title Page Footnotes:**

- We introduce a bursting tACS protocol to study semi-concurrent tACS effects in the visual system and their impact on behavior as measured by reaction time.
- Burst 10 Hz tACS (tACS_10_) applied to the visual cortex entrained γ-oscillations and increased RTs in a change-of-speed detection visual task more than 70 Hz tACS (tACS_70_) or Control conditions.
- Burst tACS_10_ also decreased amplitude of the P300 peak, while increasing α-power and γ-LZW complexity.
- Physiological and behavioral impact of occipital tACS_10_ and tACS_70_ was frequency-specific. tACS_70_ reduced γ-oscillations after 20min of tACS stimulation.
- Cognitive task may determine cortical excitation levels as measured by complexity metrics, as lower γ-LZW complexity correlates to faster reaction times.

## INTRODUCTION

A wide range of cognitive functions are mediated by the dynamic modulation of oscillatory activity within and between different brain regions. The alpha rhythm (8 - 13 Hz) has been associated to top - down processing [5, 6], mediating attentional processes [7] and linked to functions such as working memory or visual perception. High frequency gamma rhythms (30 - 80 Hz) have been associated with feature binding [8], learning or attention [3]. Although such electrical field potentials reflect the balance of synaptic excitation and inhibition [9, 10], a clear association between cognitive process and brain dynamics remains unresolved. In particular, although correlations between cognitive processes and oscillation features have been revealed, causality remains an open question [11].

Dynamic structures of brain activity, such as correlations over many timescales or self - similarity, can be captured using nonlinear or algorithmic complexity estimation methods, such as the Lempel – Ziv - Welch complexity algorithm (LZW) [12]. The LZW algorithm provides an upper bound to the algorithmic or Kolmogorov algorithmic complexity exploiting repeating patterns in the data, and summarizes it using the underlying repeating string patterns [13, 14]. Whether we adhere to an algorithmic information definition of complexity, or even if we use more classical entropic or fractal ones, we can ask what is the relationship of such metrics (derived from brain data) to behavior, health or conscious level, for example [9,10].

In turn, neuromodulatory transcranial electrical stimulation techniques such as transcranial current stimulation (tCS, also known as tES) allow for the modulation of brain activity and behavior [15] through the generation of weak electrical fields within a physiologically relevant range [16]. Transcranial current brain stimulation encloses a family of related noninvasive techniques which include direct current (tDCS), alternating current of sinusoidal form (tACS), and random noise current stimulation (tRNS). More generally, these techniques are based on the delivery of weak currents through the scalp (with electrode current intensity to area ratios of about 0.3 - 5 A / m) at low frequencies resulting in weak electrical fields in the brain (with amplitudes of about 0.2 - 2 V / m) [17, 18]). Such electrical fields are unlikely to produce per se action potentials, but have an influence on the likelihood of neuronal firing by altering the transmembrane potential. The sub-threshold effects of tCS depend also on the temporal characteristics of the electrical stimuli, as simple models suggest frequency dependent sensitivity of individual neurons to tACS (resonance). **The basic mechanism for interaction in tCS** is thought to be the coupling of electrical fields to elongated neuronal populations such as pyramidal cells, where sub - threshold polarization occurs at the soma and along fibers. Physically, the external electrical field forces the displacement of intracellular ions (which move to cancel the intracellular electrical field), altering the neuron's internal charge distribution and modifying the transmembrane potential difference, which in turn affects firing rates and derived plastic effects. Realistic biophysical models of current propagation can be used to estimate [18] and optimize [19] the electrical fields generated by multichannel tCS montages.

Our motivation in this work is the vision that non - invasive tCS can be used to probe causal relations to link physiology with behavior, and that the immediate physiological effects of tCS as measured with EEG offer an unexplored window to study such relationships [20]. In the longer term, we believe that elucidating the interaction between non - invasive brain stimulation and endogenous brain activity - and behavior - will enable the development of individualized adaptive stimulation paradigms for clinical applications. In this study we aim to provide new evidence for frequency dependence of tACS effects in the visual cortex and further discern the functional role of oscillations in cognition.

While a multitude of studies address the clinical implications of these techniques [21, 22], tCS provides also an opportunity to probe the causal role of brain oscillations in behavior. Several studies report behavioral enhancement after the application of tACS in processes like working memory [23] contrast sensitivity [3, 24, 25] or fluid intelligence [15]. Physiologically, behavioral changes after tACS have been associated to an **enhancement** of same - frequency oscillatory power. In hand with this enhancement of neuronal activity, tACS is thought to **entrain** spontaneous brain oscillations in the range of the stimulation frequency [26], as has been observed in - vitro [27]. However, the enhancement and entrainment of brain oscillations in humans appears to be state - dependent [27 – 29]. In fact, differences in endogenous oscillations alter the behavioral and electroencephalographic (EEG) effects of tACS as measured in the motor [27, 30], prefrontal [15, 31], and parieto - occipital cortex [32, 33], where the response of brain activity is seen to depend **non - linearly** on the precise relationship between endogenous and exogenous dominant frequencies. In particular, the presence of entrainment and resonance depends on the dominant oscillatory activity of the network, as proposed by in - silico models, where entrainment can be observed in 10 Hz - oscillating for tACS frequencies lower than 50 Hz [34]. In parallel, animal studies show that networks with pronounced endogenous activity impede the entrainment of activity patterns in - vitro within the tested frequency range (0.5 – 2 Hz) [27].

Here we aimed to test the relationship between observed visual cortex oscillations and behavior in a visual task, exploring in turn to what extent tACS can be used to enhance, entrain or otherwise modify endogenous oscillatory activity in healthy adults. To achieve this, we measured the behavioral and physiological impact of tACS in the occipital cortex through EEG while participants conducted a visual change – of - speed detection paradigm (see Figure 1A) as already used by other authors [4]. The task induces a reliable increase of γ - activity (50 - 80 Hz) in the early visual cortex, as well as a reduction of α (8 - 13 Hz) and β (13-25 Hz) oscillations at both visual stimulus onset (VSO) and at the change – of - speed onset (**CSO**) [35, 36]. **Crucially**, trials with faster response are associated with stronger γ - power [4], which establishes a clear if non - causal relationship between physiology (γ - power) and behavior. With the objective of **interacting** with the specific dominant frequencies at particular intervals of the task [4, 36], we designed a burst - tACS protocol that stimulates the occipital cortex during 5s after VSO at 10 or 70 Hz frequencies (tACS _70_ and tACS _10_ respectively, see Figure 1B and 1C).

**Figure 1.**
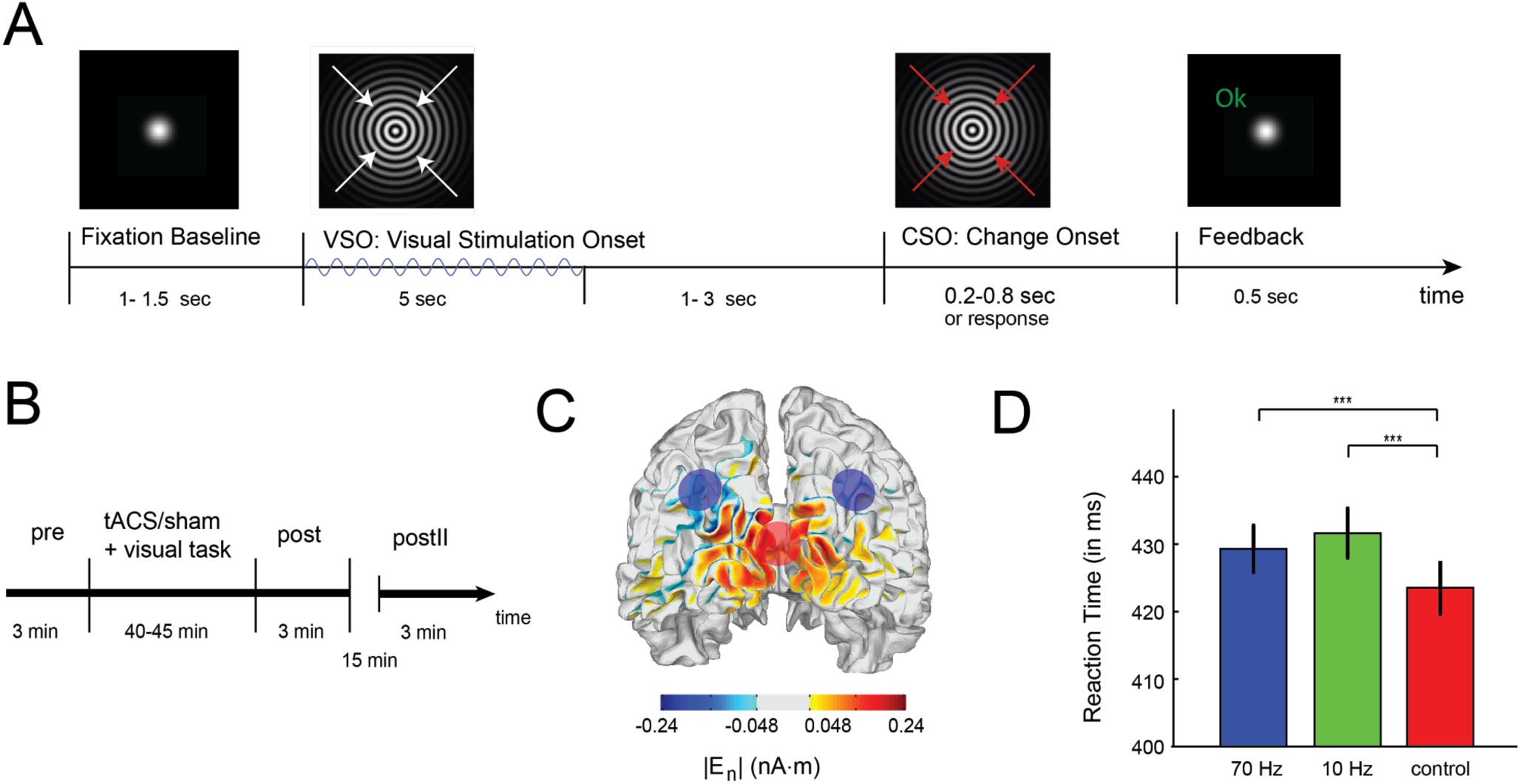
Experimental paradigm and behavioral responses. (A) Visual change - detection task. Subjects were instructed to report the change of speed of an unpredictable visual stimulus. The fixation period was followed by the presentation of a sine - wave grating moving inwards, centered at the fixation point. Bursts of tACS are delivered concurrently at the onset of visual stimulation (VSO) for 5 s in tACS sessions. At a random time between 1 and 3 s, the velocity of the moving grating increases (CSO). Subjects reported change of speed by a keypress with right finger and received feedback OK / KO on correct detection (less than 0.8s after CSO). (B) Experimental procedure. Each session starts and ends with 3 minutes of eyes open resting state (pre, post and postII) during fixation. tACS / control intervals contained 240 trials of the visual task, adding up to 30min of stimulation. (C) Current flow of the tACS using a multi - electrode optimized montage [19], revealing highest current flow in occipital cortex (stimulation electrodes located at PO3, PO4, Oz). (D) Reaction time in ms for the different stimulation protocols.

To characterize tCS derived changes in brain dynamics, we propose the use of algorithmic complexity estimation metrics. To the best of our knowledge there are no studies on algorithmic complexity - as estimated by LZW - of brain oscillations under tACS. However, various authors have studied this metric in other scenarios. For instance, stroke patients, schizophrenia, and depression patients display higher LZW complexity on both a spontaneous and a cognitive task - related EEG activity compared to age - matched healthy controls (e.g., [37]). Spontaneous EEG complexity seems to decrease during anesthesia and NREM sleep. Casali et al. [38] found it decreased also in patients with Unresponsive Wakefulness Syndrome (UWS), Minimally Conscious State (MCS) or Emergence from MCS (EMCS), and in Locked - in syndrome (LIS). Also, complexity during mental arithmetic seems to decrease compared to rest in schizophrenia, depression and in healthy controls [37]. Some of these findings are also supported by MEG (magnetoencephalography) studies. MEG signals from schizophrenic patients seem to have higher LZW complexity compared to healthy controls [39], and depressed patients seemed to have higher MEG pre - treatment complexity that decreases after 6 months of pharmacological treatment [40]. Although MEG and EEG measure brain activity differently, it seems that their underlying complexity patterns follow a similar behavior. Other MEG studies have revealed decreased complexity in MEG signals in Alzheimer's patients compared to age - matched healthy controls [41], increasing complexity until the sixth decade of life in healthy subjects, and decreasing complexity after this age, as well as higher complexity in females compared to males [42]. A recent MEG study showed that LZW complexity increases during a psychedelic state of consciousness induced using ketamine, LSD, and psilocybin compared to a placebo effect [43].

Based on previous studies, we reasoned that tACS _70_ and tACS _10_ would have a differential impact on brain dynamics. First, we expected that tACS _70_ and tACS _10_ would interact with oscillatory activity at the visual cortex at those frequencies, reducing reaction times when enhancing γ oscillations. Second, that entrainment, defined as end of stimulation phase-locked oscillations, should be observed after tACS bursts. Finally, that tACS _70_ and tACS _10_ would have a differential impact on brain complexity during stimulation.

## RESULTS

In the results and posterior analysis we focus on the EEG bands and electrodes of interest: Alpha and Gamma at posterior electrodes, where the stimulation was targeted.

### (1) Reproducing the neural signature of the task with Control sessions

The visual change - detection task has been linked to the activation of early visual areas, where it induces a reliable increase of γ - activity and reductions of alpha and beta oscillations at both the onset of visual stimulation (VSO) and at CSO (see Figure 1A) [4, 35, 36]. Consistent with previous reports, in Control sessions we observed a γ power enhancement in occipital electrodes (Figure 2C) at 0.2 seconds after CSO that decays around 0.4s. Alpha and Beta power display sustained reduction for longer latencies, co - occurring with the event - related potential (ERP) associated to CSO (Figure 2A). Taken together, these findings replicate the well - known neural signature associated with the processing of visual stimuli [4, 35, 36].

**Figure 2.**
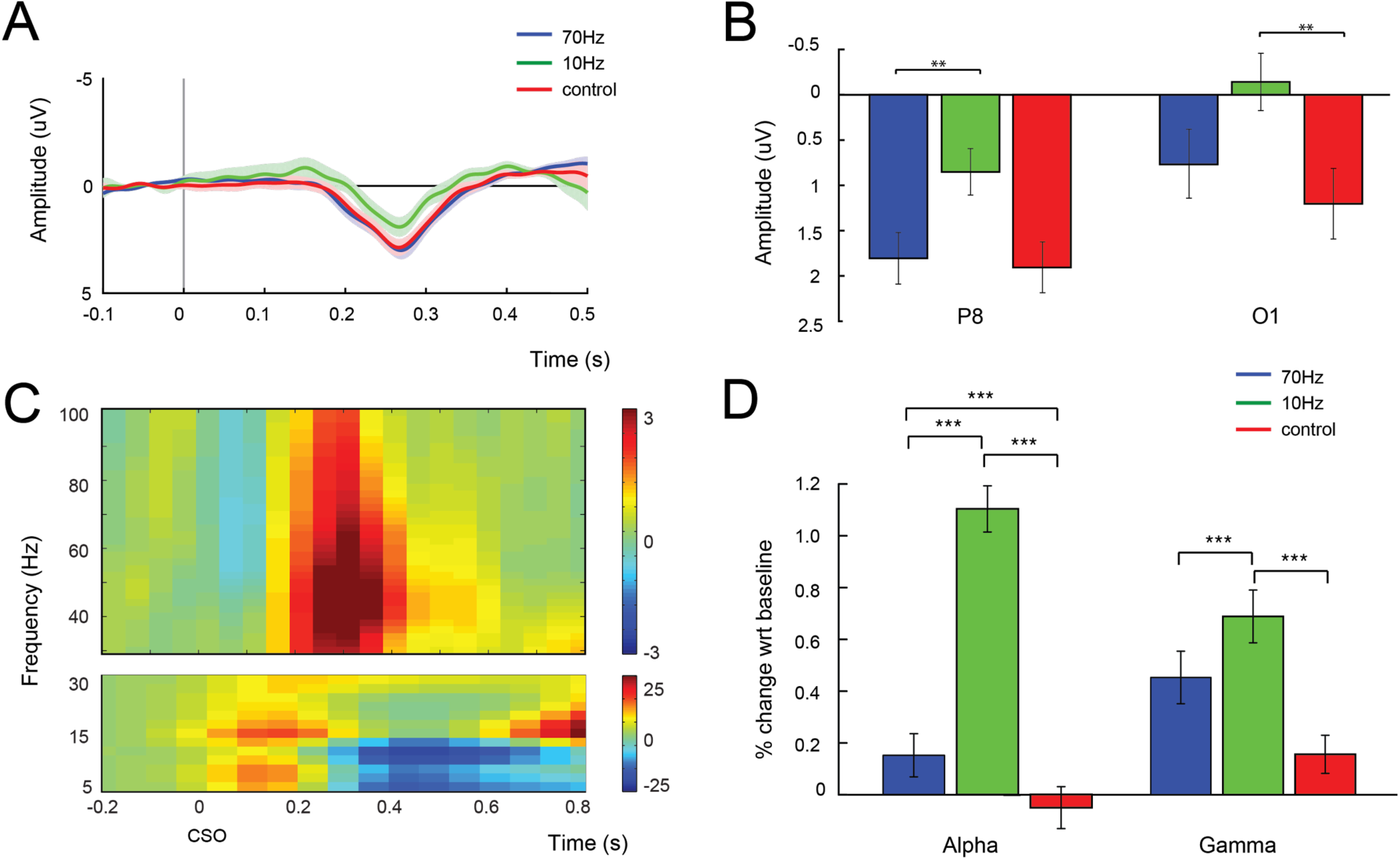
Visually induced responses in the EEG signals in time and power. (A) Event - related potential (ERP) associated to the CSO (baseline [ - 200 0] ms) for the different stimulation protocols at electrode P8 and (B) Average P300 amplitude at P8 and O1 electrodes. (C) TFR of the control sessions, expressed as a percentage of change with respect to baseline [ - 200 0]. (D) Percentage of change of incoherent α and γ power immediately after tACS bursts in occipital electrodes (0.25 - 1s after burst at O1O2). Asterisks in A indicate statistical significance as assessed by pairwise analysis of GLMM factors (p < 0.001).

### (2) tACS impacts CSO event related potentials

Event Related Potentials (ERP) are widely used in cognitive studies as their peak amplitude is thought to reflect an alteration of cortical excitability reflecting a temporal realignment of neuronal activity to stimuli [44]. In particular, larger ERP peak amplitude reflects a greater temporal alignment of neural activity with a particular stimulus, and higher excitability [10, 45]. The P300 component is produced by a distributed network of brain processes associated with stimulus - driven attention and memory operations and its amplitude and latency change as a function of cognitive resource allocation [44]. In this study, we examined the ERP that arises at the onset of change – of - speed (CSO) to test whether tACS interferes with visual excitability in a frequency - specific fashion (Figure 1A). We extracted the amplitude of the P300 associated to CSO (Figure 2A) and observed that in occipital electrodes the amplitude of the P300 is reduced with tACS _10_ stimulation as compared to both tACS _70_ (p < 0.05) and control sessions (p < 0.01) (Figure 2B) in the left - occipital cortex (electrode O1). Interestingly, this decrease in P300 amplitude in tACS _10_ sessions is also observed at P8, suggesting a possible alteration of the dorsal attention network.

### (3) tACS impacts endogenous oscillatory activity power (incoherent power analysis)

To further study tACS modulation of ongoing activity, we examined whether tACS alters α and γ power a 1s window shortly after each stimulation burst (**tCS condition**) and compared that to baseline (pre - EEG intervals, see Figure 1B). We extracted the power of oscillatory activity through the FFT, binned the frequency spectrum into bands of interest (α = [8, 13] and γ = [60 - 80] Hz), and tested whether the tACS protocols modulated power at these bands in occipital electrodes through a Generalized Linear Mixed Model (GLMM, see Supporting Information for details). The GLMM revealed a significant interaction between tACS protocol and EEG - interval (p < 0.001, t - value = 10.663). Further pairwise analysis of the model factors revealed a significant increase in α-power at occipital electrodes after tACS _10_ as compared to tACS _70_, while γ-power in occipital electrodes after tACS _70_ was **lower** as compared to tACS _10_ (all comparisons p < 0.001, see Figure 3B). Moreover, no significant change in γ-power in occipital electrodes after tACS _70_ in the **post conditions** was observed compared to Control sessions or to pre - EEG. Instead, γ - power seemed to decrease, as the stimulation blocks were finishing up (Figure 4A), hinting to a possible accumulative effect of tACS especially important at high stimulation frequencies.

**Figure 3.**
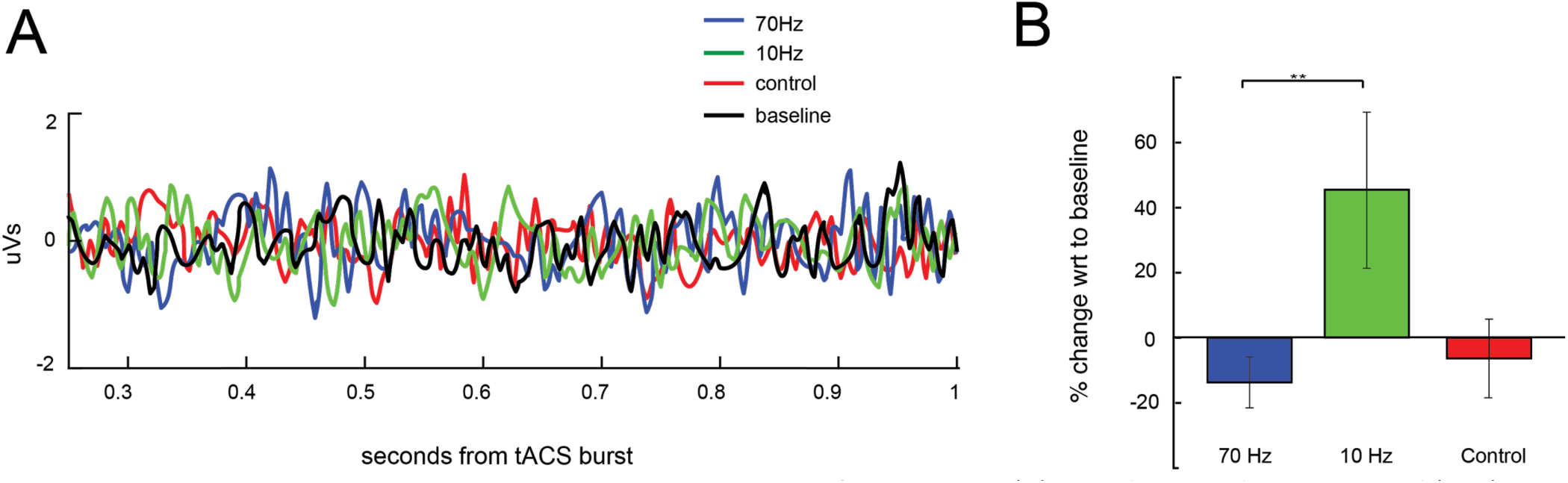
tACS Stimulation Related Potential and power change of EEG signals. (A) Stimulation Related Potential (SRP) associated to the tACS end – of - burst for a representative subject. Baseline is taken from the pre - EEG interval, see Figure 1A). (B) Percent of change in phase - locked γ power at the onset of tACS bursts in occipital electrodes (0.25 - 1.25s after tACS burst at O1O2).

**Figure 4.**
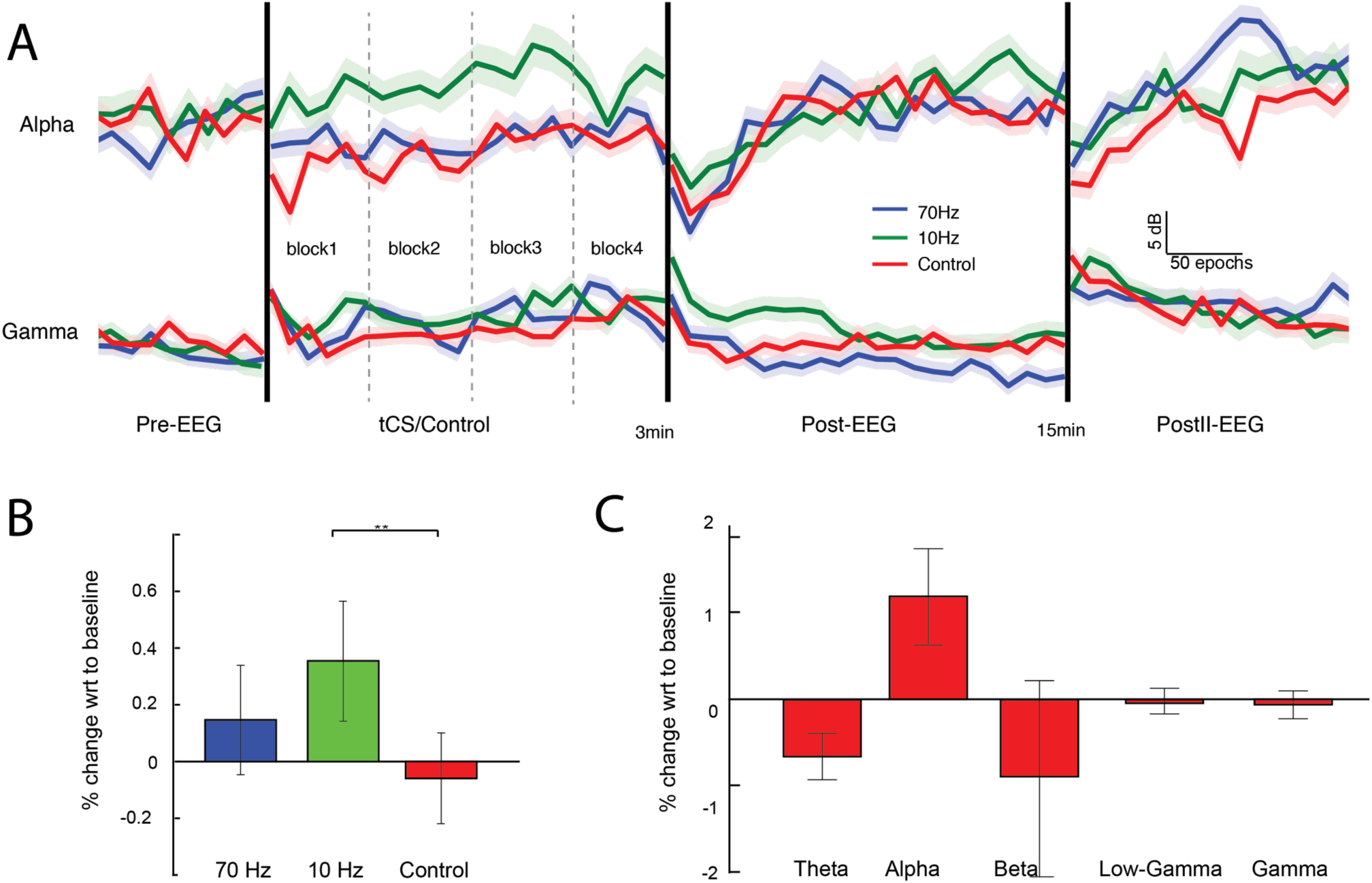
tACS after effects in power and signal complexity. A) Power change over time for 1 - s epochs across for each experimental session. B) Change in LZW at γ frequency at during the tCS condition at occipital electrodes. C) Complexity signature of the behavioral task in Control sessions.

One potential concern is that the observed power change due to tACS _10_ may to be due to an accumulative effect (plasticity) of tACS and not due to entrainment. To test for a possible interaction, we defined a GLMM where trial number is a fixed effect and found that the described effects are present in all 4 blocks of the task (p < 0.001).

Another potential concern is that this effect is not spatially localized to the tACS stimulation site (Figure 1C). To assess this, we defined a GLMM for parietal electrodes (P7 and P8). No statistical differences for α - power across tACS protocols (control included) were found. Interestingly, γ - power after tACS _70_ in parietal electrodes remained lower than γ - power after tACS _10_ (p < 0.001), as observed in occipital electrodes. This suggests that, while the modulation of α - power was spatially localized at the stimulation electrodes, modulation of γ - power recruited other cortical areas out of the focus of stimulation.

To further investigate to what extent tACS modulates oscillatory activity in a lasting manner, we analyzed the EEG at **post - EEG** intervals (3min resting state intervals after the end of the behavioral task, see Figure 1B) and 15 min after that (postII EEG intervals). We defined a GLMM where tACS stimulation type and EEG - intervals are treated as fixed effects and explored their interaction (see model details in Supporting Information). The interaction between tACS stimulation type and EEG - interval was confirmed by a pairwise analysis of the GLMM factors, all of them significant with p < 0.001.

Interestingly, we found that the α and γ - power alterations observed after tACS bursts in the tCS condition were maintained in both post and postII - EEG conditions (Figure 4A). In particular, the α - power changes observed during tACS _10_ were also observed in post and postII - EEG intervals (as compared to control and tACS _70_, p < 0.001), replicating previous results reported in both bursting [2, 46] and continuous α - tACS protocols [27, 33, 47]. Similarly, the γ - power increase observed during tACS _10_ stimulation was also observed in post and postII - EEG intervals (as compared to tACS _70_ and control, p < 0.001). Occipital γ - power in tACS _70_ post sessions was statistically smaller than the γ - power observed in Control sessions. Pairwise analysis of GLMM factors revealed that occipital γ - power during tACS / control sessions was larger than γ - power at post and postII intervals, and that α - power during tACS / control sessions was smaller than α - power at post and postII intervals), revealing the expected modulation of oscillatory activity by the task [35].

### (4) tACS impacts locked responses in time and frequency (coherent power analysis)

In order to test whether tACS produces an entrainment effect on endogenous oscillations, we examined the responses of brain oscillations phase - locked to tACS burst in terms of its Stimulation Related Potential (SRP), the evoked potential that is expected to occur due to tCS. The tACS bursts were configured to end a zero phase (zero current). This time point defined the origin for stimulation phase locked EEG analysis, giving rise to Stimulation Related Potentials akin to ERPs.

SRP responses at occipital electrodes for all three stimulation protocols are displayed in Figure 3A (between 0.25 and 1 s after tACS burst). Although no clear oscillations at the tACS stimulation frequencies are visible, we tested whether the oscillatory response phased-locked to the tACS burst (and thus, coherent) is modulated by the presence of tACS. For that, we extracted the complex FFT of oscillatory activity from the SRPs at occipital electrodes, binned the frequency spectrum into bands of interest (α = [8, 13] and γ = [60 - 80] Hz), averaged it across epochs, and finally computed its modulus. Gamma band tACS phase locked activity was **reduced** compared to baseline in tACS _70_ sessions (t - test, p = 0.096, Figure 3B), but **increased** in tACS _10_ sessions (t - test, p = 0.076) – both at trend level. In the control sessions, no change in the coherent γ - band power was observed (t - test, p = 0.606). Such changes in tACS - locked γ - power seem to **increase** further in tACS _10_ as compared to tACS _70_ (rankusm u - test, p < 0.01, Figure 3B) in occipital electrodes. No differences in phase - locked α - band are observed in any of the sessions.

### (5) tACS and LZW complexity

As previously discussed, algorithmic complexity provides the means to study the structure of oscillatory brain dynamics beyond stationary methods based on spectral features. To further elucidate the manner in which tACS affects brain complexity, we estimated the LZW metric for the tACS _10_, tACS _70_ and control conditions using the EEG data after each tACS burst. The results are presented in Figure 4B, which shows an increase in LZW complexity during tACS _10_ compared to the control condition in the high γ - band (p < 0.05).

### (6) Behavioral responses are altered during tACS

During the visual task subjects were instructed to report the acceleration of an inward - moving grating (Figure 1A). We measured behavior through the analysis of the Reaction Time (RT) and the Percentage of Correct Responses (PC). The interaction between session and behavioral metrics was assessed with GLMM, where both subject and trial number were defined as random factors (see Supplemental Information). RTs of trials where tACS was active were significantly longer than observed in control trials (p < 0.001), as shown in Figure 1D. The slowing due to stimulation was observed in all the blocks of the task (interaction of tACS stimulation type and RT is significant in all the blocks, p < 0.001), and is not tACS - frequency specific (p = 0.1741). Percentage of correct responses (PCcontrol = 97.6 (± 0.4), PC _10Hz_= 96.8 (± 0.5), PC _70Hz_= 96.9 (± 0.7), mean ± ste) was not altered by the presence of tACS stimulation (p > 0.01). Note that, while RT seems to be altered by the presence of tACS, these differences cannot be uniquely attributed to a physiological impact of the tACS, as the sensory reports of participants exposed to active vs. control tACS sessions were different. In particular, 66% of participants reported feeling no stimulation in control sessions, in contrast to the 16% and 26% of participants who reported no - stimulation in tACS _10_ and tACS _70_ sessions respectively. Thus, control sessions cannot be considered sham (see Materials and Methods).

### (8) Predicting behavior from oscillatory activity

To further understand the relationship between reaction time and its neural signature, we tested for the ability of oscillatory activity to predict behavioral responses (RT) as described in [4], where γ-band activity in the 50 – 80 Hz range of the calcarine sulcus was found to predict short reaction times in the particular visual change - detection task. This relationship was replicated in our control sessions, where the Pearson correlation between the γ - power and the RT displayed a significant correlation in occipital electrodes (r = - 0.05, p < 0.05 two - tailed one - sample t - test). Next, we proceed to repeat the correlation analysis with tACS _70_ and tACS _10_ sessions. As in control sessions, γ - power correlated with slower RT, maintaining the physiological relationship between γ - power and behavior. Interestingly, in tACS _10_ sessions, trials with higher power in low - γ frequencies (30 – 40 Hz) at occipital electrodes predicted shorter RT, extending the frequency band that correlates with behavior reported in [35].

Regarding complexity metric as predictors of behavioral responses, there is a significant positive correlation between LZW complexity in the γ - band and RT in control sessions, indicating that complexity in occipital cortex decreases with faster responses (p < 0.05 two - tailed one - sample t - test). Interestingly, tACS alters this relationship: no significant correlation appears at tACS _70_ sessions, while an increase in γ - band complexity at tACS _10_ sessions predicts slower responses (p < 0.05 two - tailed one - sample t - test).

## DISCUSSION

In this study, we examined first how tACS alters short-term physiology using a burst - EEG protocol that provides a small delay window to study concurrent tCS effects, and then how such effects are causally related to both secondary physiological effects and behavior, a technique already exploited with other stimulation techniques such as TMS [48]. tACS was used to probe the causal relationship between oscillatory activity within the occipital cortex and change – of - speed visual task, where subjects are instructed to detect a change on the acceleration of the visual stimuli (Figure 1A). This task produces a well - known and reproducible spectral signature in the visual cortex as assessed with MEG [4, 35] and EEG [36], where it induces a reliable increase of γ - activity and reductions of alpha and beta oscillations at both VSO and CSO (see Figure 1A) [4, 35, 36]. A direct correlation with behavior was established in [4], where it was reported that γ - band within occipital cortex (60 – 80 Hz) predicts shorter reaction times. Here, we interrogated this relationship between physiology and behavior by perturbing the visual system with tACS at 10 and 70 Hz.

In control sessions, where no tACS was applied (0 currents), we reproduced the increase in γ - power with respect to baseline associated to shorter reaction times (RT), as reported in [4]. When stimulating with tACS _10_, γ - power increased even more but surprisingly this enhancement of γ activity correlated with an increase of the RTs (slower responses), reversing the relation in the conventional control condition. This antagonistic relationship between physiology and behavior was observed in both coherent (i.e., phase - locked to the tACS cycle) and incoherent power analyses. While concurrent tACS _10_ enhanced α - oscillations (8 – 13 Hz), as previously reported in literature [26, 27], in our study, tACS _10_ also enhanced and entrained γ - oscillations. In addition, the concurrent α - power increase due to tACS _10_ in occipital electrodes was also observed in post and postII - EEEG intervals (as compared to control and tACS _70_, p < 0.001), replicating previous results reported in both bursting [2, 46] and continuous α - tACS protocols [27, 33, 49].

On the other hand, tACS _70_ stimulation had no statistically significant impact on concurrent γ - oscillations (neither incoherent or coherent power), while RT increased at the trend level. Surprisingly, tACS _70_ reduced γ - power at post - EEG intervals, after the task and tACS stimulation had finished. The same trend was observed at the end of experimental blocks (60 trials), where γ - power seemed to decrease in tACS _70_ sessions (trend). This relationship appears to support the established relationship between physiology and behavior (at the trend level), where lower γ - power correlates with longer RT.

During control sessions, a decrease in γ - LZW (with respect to baseline) predicted shorter RTs. Stimulating with tACS _10_ increased γ - LZW, and the increase was associated to longer RTs. Our analysis establishes a relationship between a complexity metric and behavior that complements others observed in literature, where it is reported that LZW decreases in schizophrenia, depression and in healthy controls when the participants perform a mental arithmetic task compared to their resting state EEG [37]. While the precise neurophysiological interpretation of the complexity metrics needs further work, previous studies suggest that LZW is a non - linear estimator of cortical excitation [50]. In fact, the presence of the task itself without tACS (Control condition) already induced a change in brain complexity as compared to resting pre - EEG (Figure 4C), suggesting that the presence of a cognitive task induces a change in the cortical dynamics that can be estimated through the analysis of complexity. A possible explanation for this relationship may be that when responses are fast (and correct) cortical circuits are more engaged in the task, and this restricts their dynamics to a particular reduced set of patterns, leading to a decrease in LZW. Since typically low complexity is related to more structure and predictability, it is expected to decrease when the brain dynamics are somehow synchronized with the repetitive pattern of the task. As such, tACS _10_ seems to make the visual system less structured and predictable, slowing task execution.

Regarding the ERP analysis, we found a decrease in ERP amplitude in tACS _10_ sessions, which are associated to longer RT, according to what we would expect in literature. Indeed, ERP amplitude has been largely used as an index of cortical activation or excitability [10, 44] as peak amplitude of the different components of ERP are affected by a shift of attentional demands or perceptual load [51], and in particular, larger P300 amplitudes are associated with faster reaction times [52]. Modulations of ERP amplitude have been reported after DC electrical stimulation, an effect that is dependent on the stimulation polarity and duration [53]. In particular, a relationship with the efficacy of stimulation was established in [54] where a decrease in ERP amplitude for tDCS responders is reported. In relation to the change – of - speed detection task, our results show that burst tACS _10_ sessions decrease amplitude of the P300 peak, while increasing α - power, γ - LZW complexity and RT, altogether indicating an underlying neural network misalignment to the stimulus or reduced activation.

Such relationship hint to the possibility that tACS _10_ increases inhibition in the visual cortex by enhancing α oscillations, ultimately reflecting a reduction of network alignment to visual stimuli [44]. These results are in agreement with recent studies that report a local encoding of visual stimuli and feedforward communication with higher cortical areas that are mediated by γ - band oscillations [4, 5], while local α - band oscillations (8 – 13 Hz) are involved in inhibitory feedback control processes [55] and long - range α - oscillations that modulate feedback communication with distant areas [5, 56]. In this line, behavioral tasks that emphasize top - down control of visual system display augmented synchronization in the low - frequency bands [5, 56], while local and intra - area synchronization in higher frequency hands (γ - band at 50 – 80 Hz) is reported in tasks that mostly recruit feed - forward communication [5, 35, 57]. In contrast, changes in γ - band oscillations induced by tACS _10_ (both phase - locked to the tACS cycle or incoherent to tACS) do not seem to alter the relation between γ - power and behavior, as in all tACS sessions trials, higher γ power correlated with shorter reaction times. The physiological changes induced by tACS may either impact the local encoding of the visual task or produce a shift in attentional resources due to the presence of tACS itself. Such difference determines whether changes in cortical dynamics imply a direct interaction with cortical dynamics or an indirect modulation due to sensory modulations. While participants were able to identify control conditions, no sensory differences were reported for the two different stimulation frequencies (tACS _10_ or tACS _70_), as evaluated by secondary effects questionnaires. Until further studies are conducted to understand sensory differences associated to tACS, we can argue for a frequency-specific modulation of cortical dynamics due to tACS.

## CONCLUSIONS

The study of high - level cognitive processes may benefit from non - invasive transcranial alternating current stimulation (tACS) as a method to interrogate causal relationships between physiology and behavior. Future advancements will provide an improved understanding of the particular relation between endogenous and tACS - induced oscillations, allowing the modulation of the endogenous oscillations relevant to the cognitive function of interest. Such interaction, however, can take several forms (up or down - regulation of endogenous oscillations or phase - locking) and the effects on specific brain dynamical aspects are starting to be elucidated like in this study – where we report differential effects of tACS _10_ and tACS _70_ on the physiological response of the visual cortex. As to study differential impact of brain dynamics by tACS and cognitive task, we introduce algorithmic complexity estimating metrics (such as LZW). As argued elsewhere by us and others, such metrics should characterize the computational characteristics of cortical circuitry as it engages with the external world [9] and should be related to power - law behavior of recorded signals, which in turn are believed to be linked to neuronal excitation inhibition balance [10, 13]. Our vision is that complexity metrics, as LZW, will enable the characterization of the excitability patterns of cortical areas in a timescale that allows to quantify excitability changes induced by the presence of cognitive tasks.

## MATERIALS AND METHODS

### Participants

Thirty healthy subjects (mean age of 26.6 ± 4.9 years, 13 male) participated in a randomized, double - blinded study, with a within - subject design. Three randomized recording sessions (control, tACS _10_ and tACS _70_) were separated by a 1 week of washout period to avoid carryover effects [58]. Participants gave written informed consent and received compensation for their participation. No history of neurological or psychiatric disorders was reported or any other contraindication to tCS [58]. The experimental campaign was conducted at Hospital Clinic and approved by the ethics research committee before start.

### Experimental procedure

After the placement of the tACS and EEG electrodes, participants were familiarized with the behavioral task. The experiment started with the recording of 3 min of eyes - open rest EEG (while gazing at a fixation point), 3 min of eyes - closed rest EEG and continued with the recording of 240 trials of the behavioral task. The task was organized in 4 blocks of 60 trials each, with 5 - 15 min breaks across blocks so that the subjects could rest. Sessions were closed with 3 min of resting - state EEG (during fixation) and 3 min of eyes - closed rest EEG at the end of the behavioral task (post - EEG) and 15 min later (postII - EEG).

### tACS protocol and EEG recording

Transcranial alternating current stimulation (tACS) was applied via three gelled Ag / AgCl electrodes of π cm ^2^ size (*Pitrodes* used with *Starstim*, Neuroelectrics) located at PO3, PO4 and Oz, placed according to a multi - electrode montage optimized for stimulation at BA17 using a realistic head model (*Stimweaver*, [19]). In the active conditions, AC currents were applied with 1.2mA intensity (the cortically normal component of the electric field distribution is shown in Figure 1C) in 5 - sec long bursts beginning and ending a zero phase. During control sessions zero current stimulation was delivered. The stimulation device was forced to end each tACS stimulation cycle at phase 0. For this reason, we adjusted the frequencies α and γ to 10.17 Hz and 69.99 Hz frequencies respectively. In each session, a total of 240 stimulation bursts were delivered at VSO (Figure 1A), adding up to a total stimulation time of 20 minutes. All subjects reported that stimulation did not induce phosphenes. Note that the intervals between tACS bursts were varied depending on the participant's response to the cognitive task, so no phase - synchrony was enforced between tACS bursts. EEG was co-recorded in 5 electrodes located in parieto - occipital electrodes throughout the duration of the experiment (Figure 1B) with the same electrode type as tACS. EEG signals were recorded at 500 S / s (24 bit) with the same device providing tACS (*Starstim*, Neuroelectrics). Electrode impedance was kept below 10kΩ, and the electrical reference placed at the earlobe.

Participants completed a questionnaire to assess the presence of visual and skin sensations after every session [58]. Sensations reported during active tACS were a mild itching sensation (33.3% of the subjects), followed by a moderate sleepiness (25.6%) and a mild discomfort sensation under the stimulation electrodes (6.67%), in a comparable range of those reported in literature [58, 59]. A single participant reported nausea after tACS (2.2% prevalence). Similar effects were also reported during the control session, as 7.78% of participants reported a mild itching sensation, 12.2% reported moderate sleepiness and a subject mild discomfort sensation under the stimulation electrodes. Finally, the 33.3% of the participants thought they were being stimulated in the control sessions, while 84% and 74% of participants reported stimulation in tACS _10_ and tACS _70_ sessions respectively.

### Behavioral task

Participants were required to respond with a keypress to the change – of - speed of an inward - moving visual stimulus (visual change - detection) in a reaction time visual task paradigm [4, 35, 36]. Each trial began with the display of a fixation point (Gaussian of diameter 0.5°), and subjects were instructed to fixate to that position through the length of the trial. After 1 to 1.5 seconds (interval randomly chosen from a uniform distribution of 1 - 1.5 s) the fixation point was replaced by a moving grating (sine wave of 5° located at the fovea contracting towards the fixation point at a spatial frequency of 4 cycles / °, contrast 100%). After 6 to 8 seconds, its velocity increased to 2.2 deg / s until response was reported or 0.8 seconds passed (CH onset, see Figure 1A). Subjects were instructed to report the velocity increase with a button press on a keyboard, which made the moving grating disappear. Feedback was provided to participants via OK / KO signs after their response. A response earlier than 0.2 seconds after CSO was reported as KO. Stimuli were displayed on an LCD screen located at 60 cm of the subject, with a vertical refresh rate of 60 Hz.

### Data analysis

The analysis was performed using customized Matlab code (MathWorks Inc. Natick, MA, USA), EEGlab [60] and FieldTrip [61].

*Behavioral analysis:* subject responses were quantified in terms of the percentage of correct responses (i.e., correct detection of the velocity change or PC) and the reaction time (RT, i.e. delay in ms between CH onset and the keypress). Sessions whose PC was smaller than 85% were rejected. The differences in RT and PC were quantified by a generalized linear mixed - effects regression model (GLMM), see Supplemental Information.

*EEG analysis:* raw EEG was cut into epochs of interest depending on whether we would analyse time - locked responses or ongoing oscillatory response. To assess visual cortex responses, we analyzed the signal of occipital electrodes O1 and O2. All epochs were filtered as follows: epochs were individually transformed into the spectral domain using the direct FFT transformation. Bins not in the frequency ranges of [60, 80] and [5, 40] Hz were removed and subsequently returning to the temporal domain applying the inverse Fourier transform. Epochs containing signals with amplitude out of the + / - 50 uV range were rejected (high amplitude threshold) after filtering. Finally, unless indicated, EEG signals were referenced to the Pz electrode.

*Subject rejection*: After artifact correction and filtering, subjects containing less than 60 artifact - free 1s epochs (at any electrode) were rejected, as a compromise between a sufficient number of epochs and subjects for a significant statistical analysis. As result, 7 subjects out of the 30 were discarded from further analysis.

*Time - locked responses*: Two different events in the task were used to study time - locked responses. tACS-burst EEG epochs were 1s long starting 250 ms after the end of tACS stimulation (see Figure 1A). Residual amplitude clipping observed at the end of tACS bursts was corrected before the filtering step as described in Supporting Information. Subject's stimulation related responses (SRPs) was calculated averaging occipital electrodes epochs in the time domain. Also, epochs of [- 0.2, 1] s around the CSO were extracted (Figure 1A) for ERP and spectral analysis in occipital electrodes.

*Ongoing oscillatory responses:* The analysis of ongoing oscillatory responses largely followed earlier work [26, 46, 62]. Resting state intervals in pre - EEG, post EEG and postII - EEG intervals (Figure 1B), were segmented into 1s epochs. During behavioral task, (Figure 1A), 1s epochs starting 250 ms from the end of the tACS were extracted.

*Spectral analysis*: Two different methodologies were used to conduct spectral analysis. On the one hand, epochs around CSO were analyzed for non-phase-locked oscillation by means of the time-frequency representations (TFRs) as described in [35]. Briefly, TFRs of frequencies between 30 - 100 Hz were obtained using the multi - tapering method, in steps of 2.5 Hz, while TFRs of frequencies between 5 - 30 Hz were computed using a wavelet transform. The TFR were computed using a Fourier transform on 1s epoched data in 50 ms steps. A smoothing of ± 5Hz (squared function) was applied around each center frequency as reported in [4, 5]. On the other hand, SRP power (coherent power) was calculated at θ = [5, 8] Hz, α=[8, 13] Hz, β = [13, 25] Hz low - γ = [30 - 40] Hz and γ = [60 - 80] Hz bands via trapezoidal integration of the power spectral density of SRP. Calculated power is referenced (divided) to band power calculated the pre – EEG - eyes open epochs.

*Complexity analysis:* As described in [12, 13], in LZW we consider a string of characters and alphabet with A symbols (typically binary) of length n. The algorithm works by initializing the dictionary to contain all strings of length one and then it scans through the input string sequentially until it finds a string that does not belong to the dictionary, and adds it to the dictionary. This process is repeated until all input string has been scanned through. Following this process we end up with a set of words c (n) that make up the dictionary. The length of the compressed string is lengthLZW < = n (an upper bound to Kolmogorov or algorithmic complexity). The description length of the sequence encoded by LZW would have length equal to the number of phrases times the number of bits needed to identify a seen phrase plus the bits to specify a new symbol (to form a new phrase), hence (1) lengthLZW = c(n) log2 [c(n) + log2 A] ∽c(n) log2 [c(n)]. The lengthLZW is normalized by the original string length. The input string is binary and is derived by taking the median of the input time series as the threshold as it is a robust metric against outliers, assigning zeros to all values below the threshold and ones to all values above the threshold. The input time series consist of 4 - sec epochs filtered in the θ = [5, 8] Hz, α = [8, 13] Hz, β = [13, 25] Hz low - γ = [30 - 40] Hz and γ = [60 - 80] Hz bands for each electrode.

## AUTHOR CONTRIBUTIONS

DI, MC, MCam, XM conducted the data recording and MC, DI, EK and ASF did the analysis. MC and GR co - designed the protocol, coordinated the data analysis. GR and EK prepared the code for LZW analysis. GR designed the stimulation montage. JV coordinated data recording and MC, DI, EK, ASF, JV, AV and GR helped write the manuscript.

## ACKNOWLEDGMENTS

This work was partly funded by Biogen Pharma. The complexity analysis part was partly been undertaken under the umbrella of the European FET Open project Luminous. This project has received funding from the European Union's Horizon 2020 research and innovation program under grant agreement No 686764.

## Conflict of interest

GR is a co - founder, shareholder and employee of Neuroelectrics, the company that provides Starstim and Stimweaver. MC, EK, DI, XM and ASF, are employees of Starlab, the company that gave birth to Neuroelectrics in 2011. All other authors declare no potential conflicts of interest with respect to the research, authorship, or publication of this article.

